# Phage-based microbiome manipulation elucidates interactions within community context

**DOI:** 10.1101/2025.06.20.660599

**Authors:** Tomas Hessler, Dawn M. Chiniquy, Benjamin A. Adler, Jordan Hoff, Eliana Tucker, Robert Huddy, Rohan Sachdeva, Shufei Lei, Susan T. L. Harrison, Rodolphe Barrangou, Spencer Diamond, Adam M. Deutschbauer, Jillian F. Banfield

## Abstract

Testing microbial interaction hypotheses remains a major challenge in microbiome research. Bacteriophages have the potential to probe organism-organism interactions within complex microbiomes, yet are rarely used for this purpose. Here, we isolated nine narrow host range phages that replicate in several ecologically important *Variovorax* species. As *Variovorax* CL14 degrades the plant root-stunting hormone auxin, we used phages to eliminate *V.* CL14 from a rhizosphere consortium and reestablished the stunted root phenotype. We used three of the phages to deplete another *Variovorax*, SCN45, from complex communities to test correlation network-based hypotheses of thiamine interdependencies. Genome-resolved metagenomics revealed that specific taxa decreased in relative abundance following *Variovorax* depletion and could be rescued by thiamine supplementation. Thus, we confirmed thiamine production as the mechanistic basis for interdependence. These experiments lay the foundation for research that employs wildtype and engineered phages to test interaction hypotheses and for targeted microbiome manipulation.

## Introduction

Microorganisms form the foundation of food webs, and profoundly influence the health of plants, humans, and other animals on a global scale. Yet microorganisms rarely function in isolation, and instead form complex communities in which a multitude of interactions shape its structure (Pacheco et al., 2019, Ma et al., 2020, Kost et al., 2023, Clegg & Gross, 2025).

Complex microbial communities have long been studied using metagenomics (Tyson et al., 2004), metatranscriptomics (Gilbert et al., 2008, Frias-Lopez et al., 2008), metaproteomics (Ram et al., 2005), and metabolomics (Segata et al., 2013), sometimes informed by stable isotope probing (Starr et al., 2018). Although these methods may generate hypotheses about organism interdependencies, our ability to test predicted interdependencies in a community context is extremely limited and largely restricted to highly specialized techniques (Pierce and Dutton 2022).

Microbial community models can infer the sum of all interactions (Skwara et al., 2023, Quinn-Bohmann et al., 2024), but lack of species-resolved mechanistic understanding limits our ability to engineer these systems. Metabolic models can predict interactions by incorporating microbial metabolic capacities (Dukovski et al., 2021), but this approach is primarily limited to co-cultures and simple synthetic communities. This motivates the search for approaches that can elucidate organism-organism interactions in complex microbial ecosystems.

Bacteriophages offer a highly targeted way to probe microbial interactions due to their inherent bacterial specificity and potent host-killing efficiency. Bacteriophages have been predominantly used as therapeutics to remove or deplete pathogenic and antibiotic resistant microorganisms (Buttimer et al., 2022; Thanki et al., 2022; Das et al., 2015, Strathdee et al., 2023; Liao et al., 2025). Bacteriophage can also be further engineered for improved host killing and broadened host range (Selle et al., 2020; Gencay et al., 2023).

Prior work has established that phage hold enormous potential for microbiome manipulation. Carlson et al. (2023) demonstrated that the ammonium-generating capacity of fresh-water enrichments could be altered through the use of applied bacteriophage. Li et al. (2022) demonstrated the depletion of a specific *E. coli* using T7 phage can modulate host phenotypes, specifically animal behaviour. While several studies have engineered phage for increased killing efficiency (Selle et al., 2020), depletion of antibiotic resistance encoding plasmids (Bikard et al., 2014) and modification of host range (Gencay et al., 2023).

Here, we isolated multiple lytic bacteriophages specific to a collection of *Variovorax* isolates, and used them to modulate microbiomes to test several predicted microbial interactions. Our findings demonstrate that phages can be used to probe microbial interactions in the context of complex microbial communities.

## Results

### *Variovorax* phage isolation

To isolate *Variovorax* phages capable of selectively killing *Variovorax SCN45* we extracted phage viromes from six rhizosphere and compost samples (Oakland, CA) and tested for infectivity against a panel of *Variovorax* strains (Methods). We isolated nine phages that can replicate in one or more *Variovorax* species. These double-stranded DNA phages ranged in genome size from 43 to 159 Kb and with GC contents of 53% to 66%. Based on protein homology, seven of these phages (all with ∼50 kbp genomes) were grouped with other RefSeq phages (Fig. 1a) classified predominantly with *Siphoviridae* (now *Siphovirus*) and share a high degree of average nucleotide diversity (ANI) with each other (Fig. 1b). The remaining two phages clustered with *Myoviridae* and *Autographiviridae* families. Two *Siphoviridae* phages (bapic4 and V.45_iii) and the large *Myoviridae* phage (V.45_v66) showed strong lytic activity against *V.* SCN45 (Fig. 1e). V.45_v66, bapic4 and V.45_iii were shown by TEM to have *Siphovirus*-like morphology, consisting of an icosahedral head and long filamentous tails, similar to the well-known lambda phage (Fig. 1d). All nine of the isolated phages are predicted to be restricted to a lytic lifestyle based on the absence of integrase genes. There was very limited cross-reactivity of the phages between the different *Variovorax* strains (Fig. 1c) based on spot assays. Only one phage, V.45_66, was able to form plaques on multiple hosts – *V.SCN45* and *V. paradoxus DSM 645* (Fig. S1).

**Figure 1.**
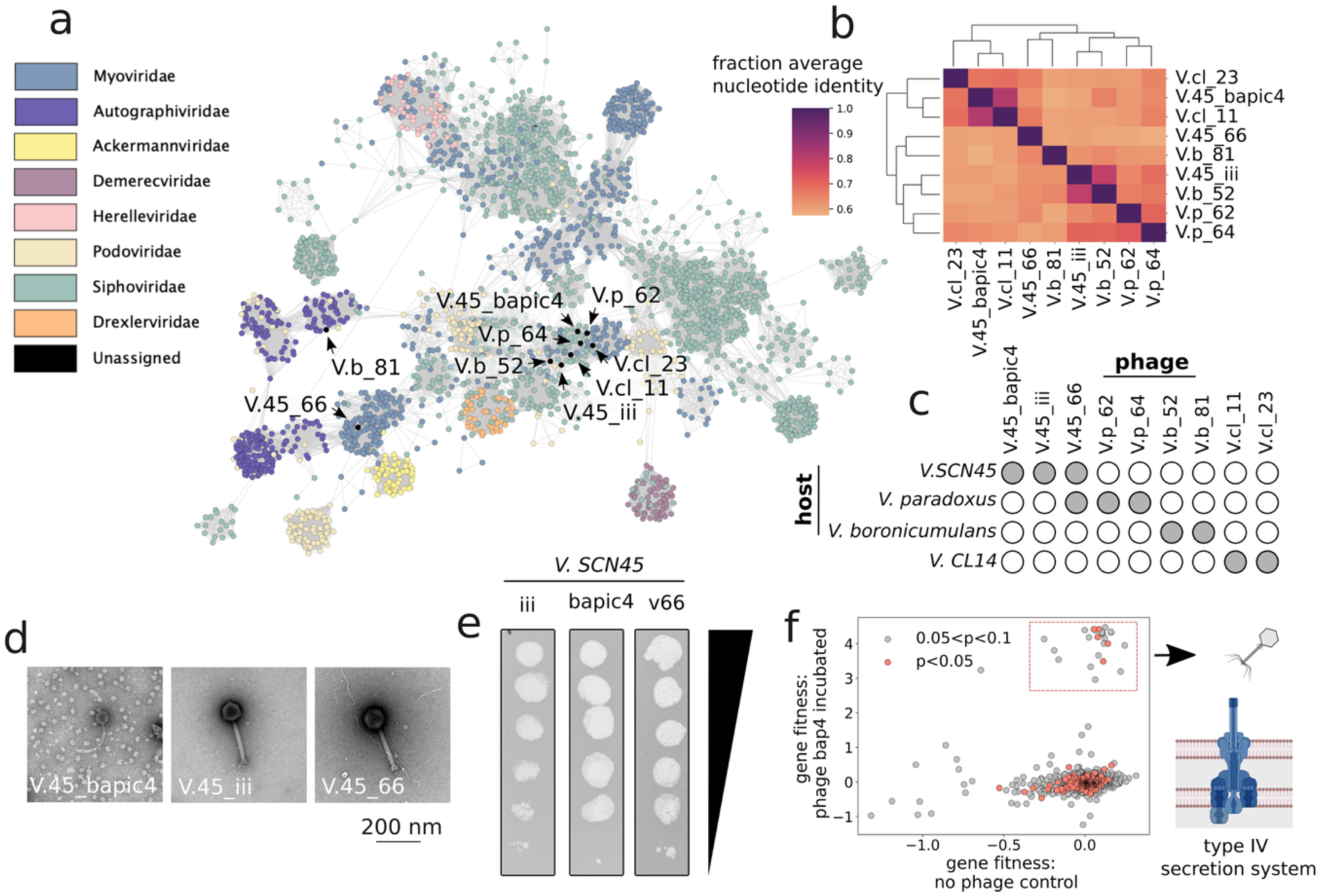
Characterization of nine isolated *Variovorax*-infecting bacteriophages. (a) Protein similarity network of phages deposited in RefSeq and those isolated in this study (black), with edges indicating a significant shared protein content. (b) Pairwise average nucleotide identity (ANI) between the nine isolated *Variovorax* phages. (c) Plaque assays demonstrate limited cross-reactivity of the nine phages across four different *Variovorax* strains, with filled circles representing the ability to plaque on a given host. These include *V.* SCN45, *V.* CL14, *V. boronicumulans* DSM 21722, *V. ginsengisoli* DSM 25185 and *V. paradoxus* DSM 645. (d) Transmission electron microscopy (TEM) of the SCN45-infecting phage, showing icosahedral heads and filamentous tails characteristic of Siphoviridae. (e) High titre phage stocks visualized by 10-fold dilution 1uL spot assays from 10^-6^ to 10^0^. (f) Genome-wide fitness profiling (RB-TnSeq) for *V. SCN45* genes, following exposure to phage bapic4, identifies the Type IV pilus as the likely receptor used by the phage for infection. Phages V.45_66 and V.45_iii generated similar gene fitness results.

### *Variovorax SCN45* phages infect via chromosome-encoded type IV pilus

To identify loss-of-function gene disruptions that could confer *Variovorax* with increased fitness under phage infection we constructed an RB-TnSeq mutant library of *Variovorax* SCN45 and employed methods similar to Mutalik et al. (2020). The disruption of genes of the type IV secretion system (T4SS) were found to substantially enhance the fitness of *Variovorax* when incubated with each of its three phages (Fig. 1f). None of these T4SS gene knockouts were enriched in the no-phage controls, indicating that the addition of phage selected for these gene mutants through the elimination of susceptible mutants.

### Evaluating Phage-Mediated Depletion in the *Arabidopsis*-*Variovorax*-*Arthrobacter* System

Finkel et al. (2020) demonstrated that *Variovorax CL14* mitigates a root stunting phenotype in *Arabidopsis* by consuming excess auxin secreted by *Arthrobacter* that leads to root stunting. To test whether such well-characterized interactions could be tested through phage-based manipulation, rather than conventional strategies, we hypothesized that a co-culture of the two organisms would result in a normal root phenotype but phage-mediated depletion of *V.* CL14 would result in the accumulation of auxin and cause a stunted root phenotype. We employed a phage cocktail of the Siphoviridae phage V.cl_11 and V.cl_23 to deplete *V.* CL14 from both pure cultures and co-cultures with *Arthrobacter CL28* inoculated onto *Arabidopsis thaliana*. Phage treatment led to the reintroduction of root stunting phenotype (Fig. 2).

**Figure 2.**
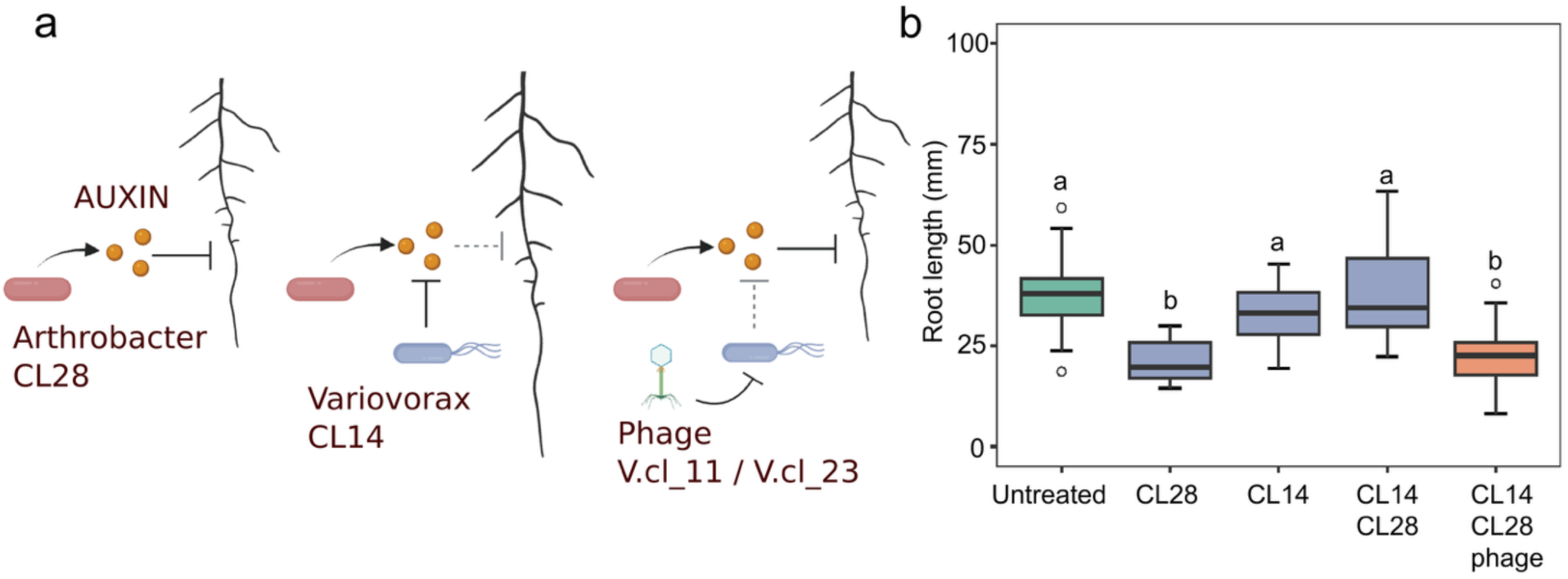
Phage-mediated depletion of *Variovorax* from an *Arabidopsis*-inoculated synthetic community induces root stunting. (a) Model of root inhibition by the auxin-secreting *Arthrobacter* CL28, which reverts to a normal root phenotype when the auxin-degrading *Variovorax* CL14 is present. If the *Variovorax*-targeting phage cocktail (V_11, V_23) is effective in the rhizosphere, this would disrupt the interaction and the root would revert to stunted growth. (b) *Arabidopsis* primary root elongation with *Arthrobacter* CL28 causes a stunted root phenotype, which recovers with *Variovorax* CL14 co-culture. The addition of the *Variovorax*-targeting phage cocktail (V.cl_11, V.cl_23) to the co-culture of *Arthrobacter* CL28 and *Variovorax* CL14 recapitulates the root stunting phenotype observed in the presence of *Arthrobacter* alone. Letters above bars indicate the results of a Tukey post hoc test (α=0.05).

### Development of stable enrichment communities

To perform community manipulation experiments it is critical to generate stable, reproducible consortia with reasonable complexity, ideally containing organisms that are predicted to be interdependent on other community members. This study leveraged inoculum from lab-scale bioreactors from which the *Variovorax SCN45* pure culture was previously isolated (Hessler et al., 2023). *Variovorax SCN45* overproduces thiamine in pure culture and was predicted to supply this essential vitamin to highly dependent thiamine auxotrophs in bioreactor communities based on ecological network analyses (Hessler et al., 2023). The thiamine auxotrophs that were correlated with *Variovorax* had very few correlations with other thiamine-producing organisms, suggesting a strong dependence.

We supplemented additional *Variovorax* SCN45 cells to increase its abundance in inoculum consortia based on previous findings that initial ratios of organisms can affect the final steady state community composition (Gao et al., 2021). The inoculum was then grown in minimal media, supplemented with various carbon sources, and passaged every 48 hours for a total of seven passages (Fig. 3a) and community compositions assessed. Four carbon sources were chosen for ongoing experiments because the communities were reproducible (Fig. 3, S2). The communities included *Leifsonia*, multiple Xanthomondales and several *Microbacterium*, taxa that were predicted to depend on *Variovorax* for thiamine (Hessler et al., 2023).

**Figure 3.**
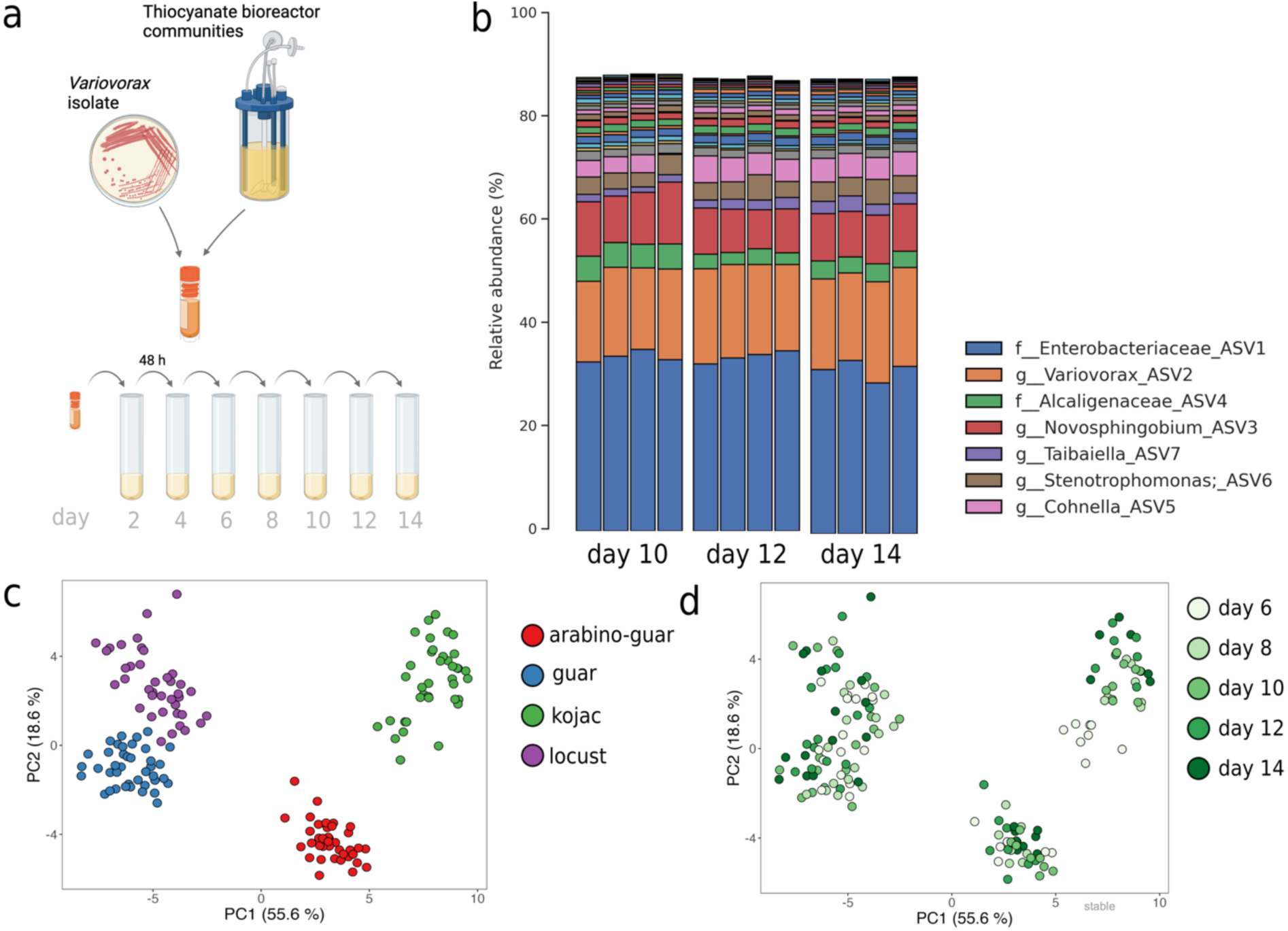
Development of stable enrichment communities. (a) Schematic of Bioreactor-derived inoculum preparation and enrichment passaging. (b) Communities supplemented with guar gum showed reproducible compositions across technical replicates and demonstrated stability across multiple passaging cycles using 16S rRNA gene amplicon sequencing. PCA was performed on robust centered log-ratio (rclr) transformed ASV relative abundances, demonstrating the compositional (c) reproducibility and (d) longitudinal stability were consistent across the cultures supplemented with arabinogalactan with guar gum, guar gum alone, kojac gum, and locust bean gum.

### Phage-mediated *Variovorax* depletion leads to broad community impacts

Following the passage of the day 8 communities, either the phage V.45_v66 or a cocktail of three *Variovorax* phages (V.45_v66, V.45_iii, and V.45_bapic4) was added to the communities (Fig. 4 a). The relative abundance of *Variovorax* in the guar gum enrichment was reduced from approximately 22% in the day 8 communities to 2.4% in the day 10 community but further recovered to 11% by day 12 (Fig. 3b). This pattern was consistent across all four communities (Fig. S2). There was minimal difference in the abundance of *Variovorax* between the single phage V.45_v66 and three-phage cocktail, except in the kojac gum-community, where the three-phage cocktail resulted in a lower *Variovorax* abundance (mean 4%) compared to the single V.45_v66 phage (mean 10%) following passage six (day 12).

**Figure 4.**
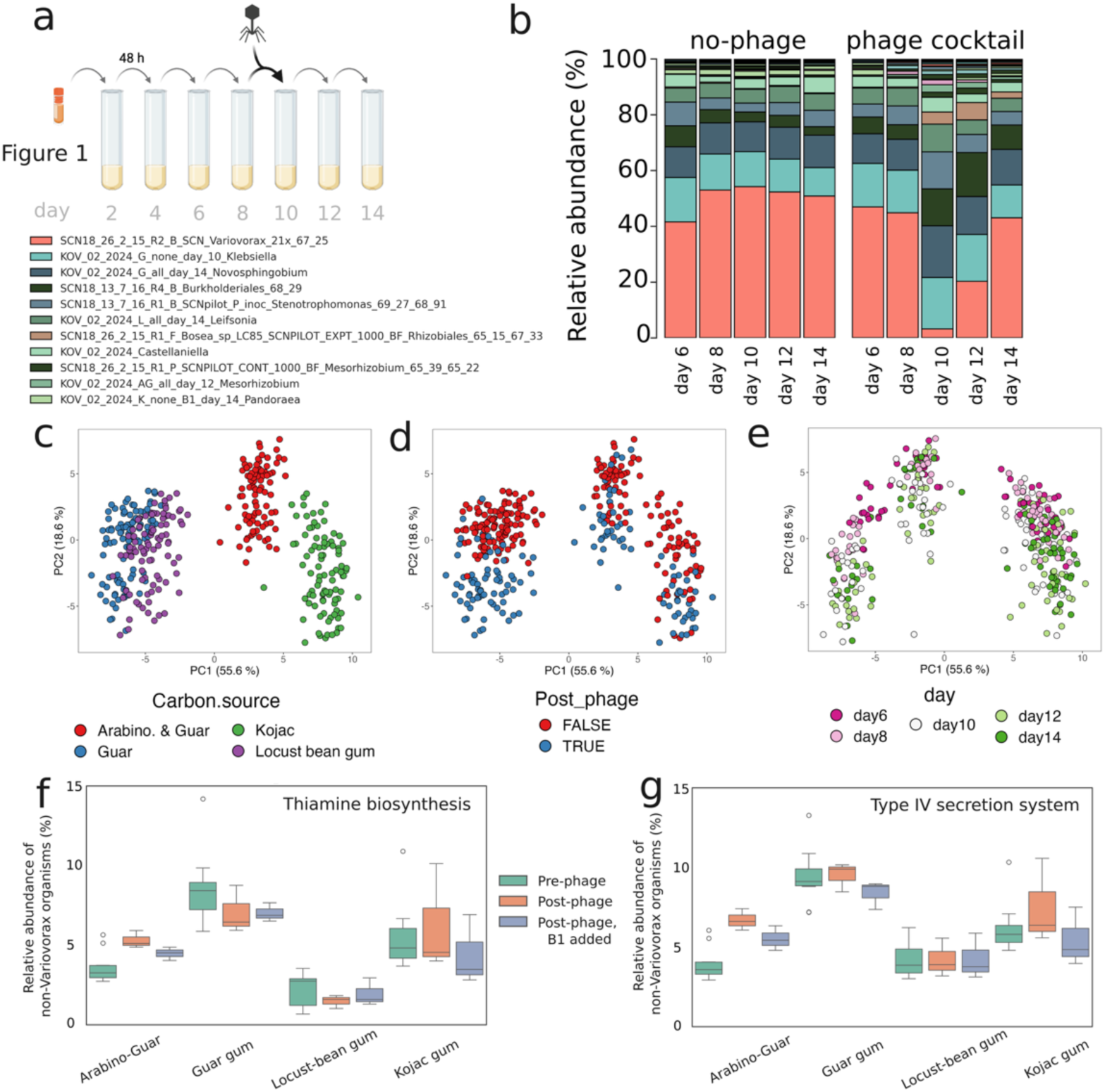
Phage-mediated *Variovorax* depletion across enrichment cultures. (a) Schematic adapted from Figure 3, illustrating the introduction of phage to 2×10^4^ plaque-forming units per mL (PFU/mL) on day 8 following passaging. (b) Genome-resolved metagenomics revealed a marked reduction of *Variovorax* (pink) on day 10 post-phage. Principal component analysis (PCA) of log-normalized abundance data from amplicon sequencing data (c) reveals a separation along PC1 driven by carbon source, while (d) PC2 captures the divergence from pre– and post-phage conditions, and (e) day of the experiment. The summed relative abundance of all organisms encoding (f) a complete thiamine biosynthesis pathway and (g) a type IV secretion system (excluding *Variovorax* from 100% relative abundance). Shown across three stages: pre-phage addition (day 6 and day 8 communities), post-phage addition (day 10, 12, and 14 communities) and post-phage communities supplemented with 10 mg/L thiamine (B1).

The overall community structure was impacted by the addition of phages, as evidenced by a notable shift in the principal component two of the PCA (Fig. 4c-e) performed on log-normalized abundance data from amplicon sequencing data. Despite phage addition, community stability was re-established, both in terms of species composition (Fig. 3c-f) and optical density over subsequent passages (Fig. S3). Bacteria other than *Variovorax* that can produce thiamine did not proliferate following *Variovorax* depletion (Fig. 4f). There was no evidence that *Variovorax* phages interacted with non-target organisms that encode a Type IV secretion system (Fig. 4g).

### *Variovorax* depletion confirms dependence of specific thiamine auxotrophs

We assessed which organisms exhibited statistically significant abundance changes between the ‘pre-phage communities’ (day six and day eight communities) and the communities following the addition of phage (day 10, 12, 14 communities). After phage administration, we added thiamine to the control enrichment communities to identify the organisms that were specifically reliant on *Variovorax* for thiamine. Amplicon sequencing of the four enrichment cultures revealed that some organisms that decreased in abundance following *Variovorax* depletion, and then increased significantly in abundance following thiamine supplementation (Fig. 5a). These included a *Lysobacter* (Xanthomonadales), a *Leifsonia,* and microorganisms belonging to the family of Microbacteriaceae (Fig. 5b). Genome-resolved metagenomics of the samples revealed that the *Leifsonia* (Fig. S5), a *Microbacterium* and a *Stenotrophomonas* (Xanthomonadales) all lack a complete thiamine biosynthesis pathway (Supplementary data).

**Figure 5.**
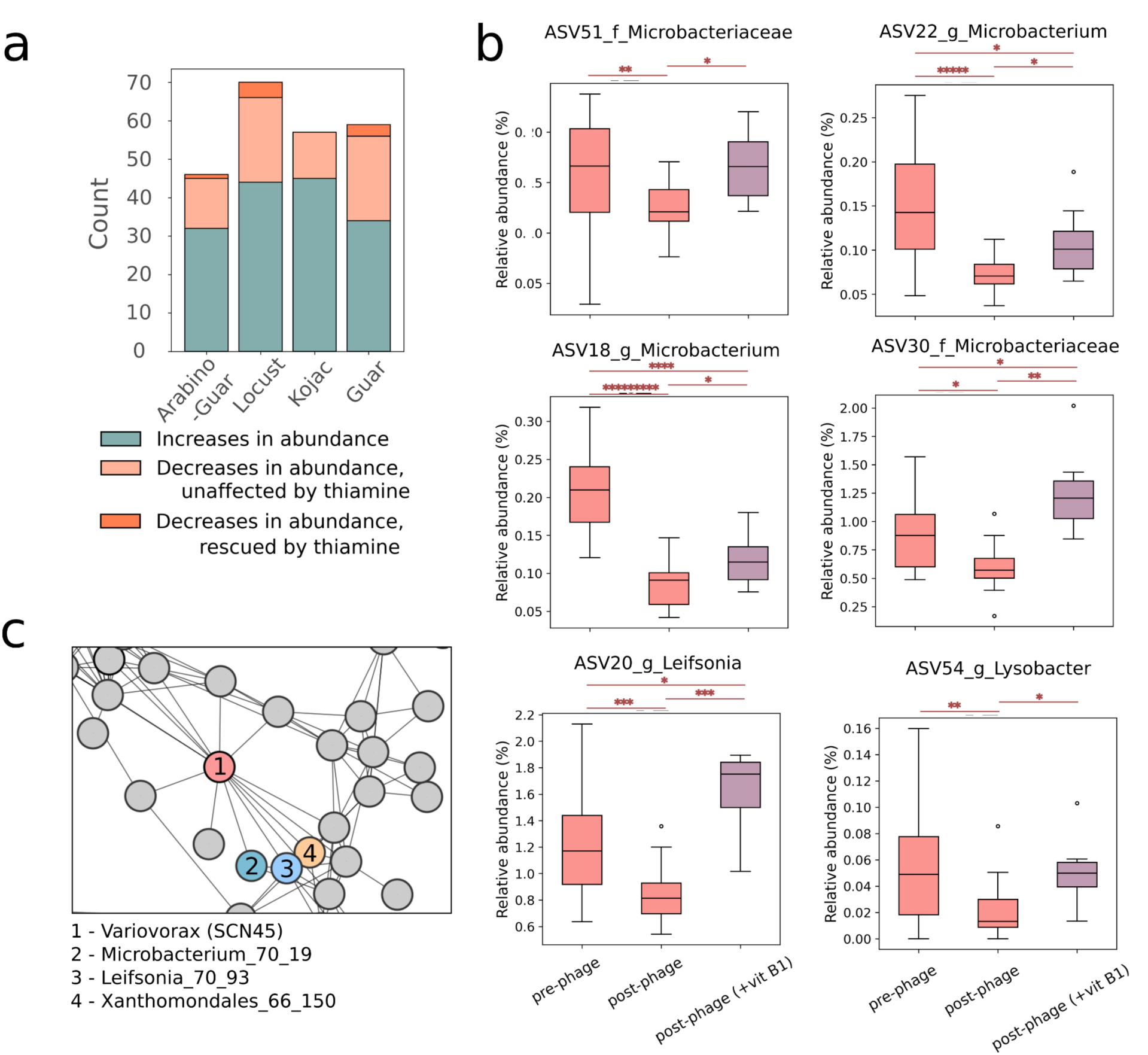
Previously identified organisms experimentally confirmed to rely on *Variovorax* for thiamine. (a) Amplicon sequence variants (ASVs) exhibiting significant changes in relative abundance between the pre– and post phage communities. Many ASVs increased in abundance following *Variovorax* depletion while others showed reduced abundance, and with a subset of these demonstrating recovery following the addition of thiamine. (b) Six ASVs corresponding to multiple *Microbacteria*, a *Leifsonia*, and a *Lysobacter* show a reduced abundance following the depletion of *Variovorax* but returned to near original levels with the supplementation of thiamine. (c) Closely related strains of these organisms appear in the correlation network that was initially used to predict *Variovorax* as a thiamine-producer and show correlations with *Variovorax* (Hessler et al., 2023).

To determine if the association between *Leifsonia* species and thiamine-producing bacteria may be more general, we assessed 32 *Leifsonia* genomes present in GTDB for the ability to synthesize thiamine. We found only a single *Leifsonia* genome to have a complete pathway (Supplementary data).

### SNPs enabled post-infection recovery of *Variovorax*

The *Variovorax* SCN45 genome is comprised of a 6.8 Mb chromosome and 1.2 Mb circular genetic element that is related to a *Variovorax paradoxus S110* sequence described as a “second chromosome” by Han et al. (2010). *Variovorax* SCN45 lacks CRISPR-Cas systems but encodes 24 PADLOC(v4.3.0)-predicted phage defence systems. We were unable to detect any deletions of these genes, or of any genes (including those of the T4SS), in post-phage *Variovorax* populations based on read mapping and gene abundances.

To seek evidence for mutations that could confer phage immunity in surviving *Variovorax* populations, we monitored *Variovorax* single-nucleotide polymorphisms (SNPs) across the four experiments. Before the introduction of phage, the majority of reads map to the reference genome without SNPs. However, following phage addition, there is a marked increase in the representation of SNPs and an increase in the number of SNP sites. In prephage communities the majority of wildtype haplotypes are present at approximately 90% (indicating that at a given SNP site, 90% of mapped reads support the wildtype base). However, following phage infection the proportion of SNP sites with the wildtype haplotype reduces, demonstrating selection of new or previously less abundant SNPs (Fig. 6a, S6).

**Figure 6.**
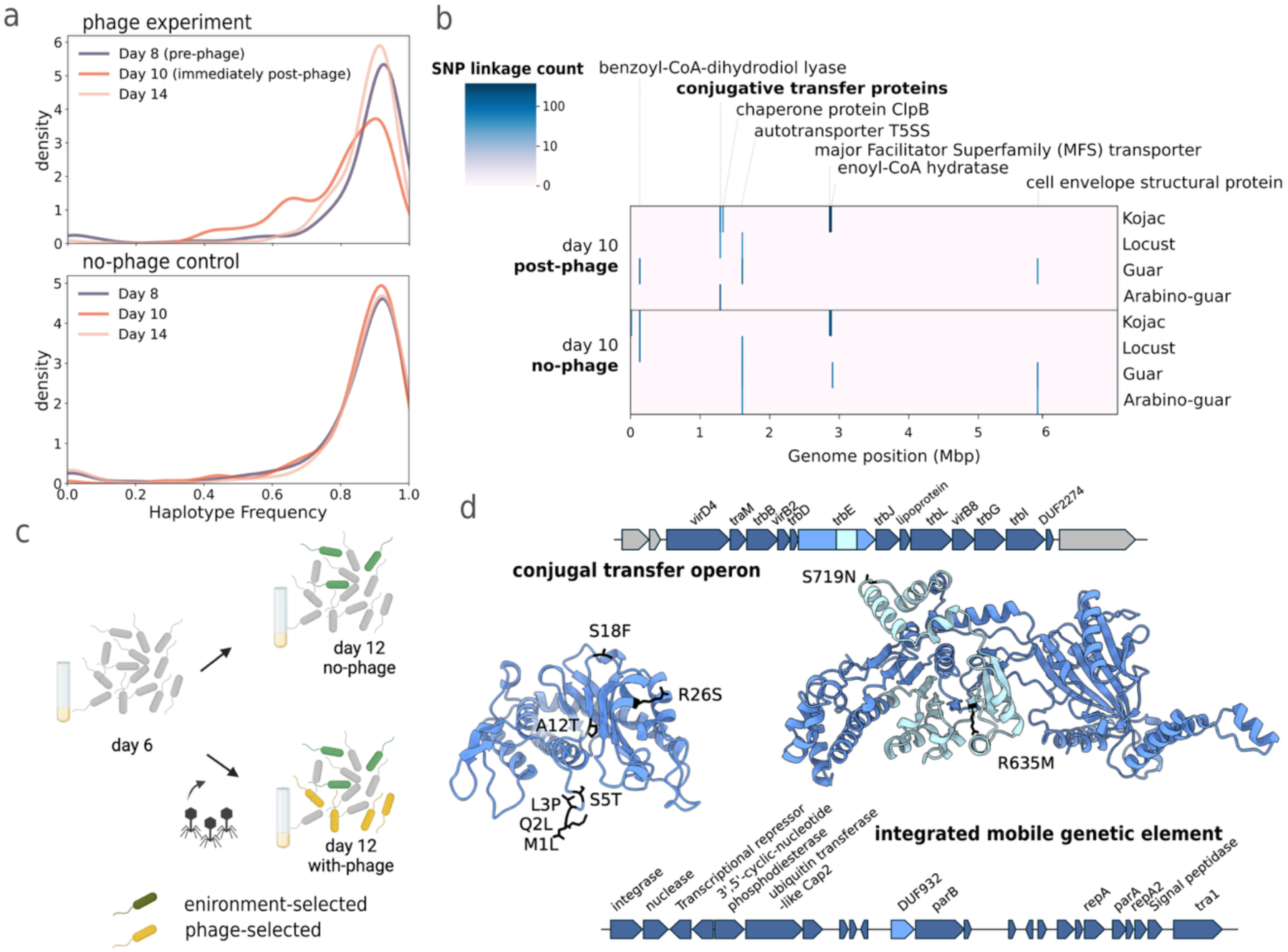
Phage infection induces shifts in *Variovorax* populations and selects for consistent gene variants. (a) Haplotype frequency distributions of *Variovorax* populations in the locust bean gum communities shows the impact by phage application, with expansion of low abundance SNPs, followed by return to pre-phage distributions. (b) Post-phage *Variovorax* populations show increased linked-SNP density in a conjugative transfer gene operon, visualised by a heatmap. Linked-SNPs associated with three 16S rRNA genes have been removed for clarity. (c) A schematic illustrating different categories for selection in day 12 communities. (d) Gene operons and AlphaFold-predicted protein structures showing consistent mutation sites within the conjugal transfer protein TrbE and domain of unknown function within an integrated genetic element.

Although most SNPs post-phage infection were synonymous, a subset was not. To identify new gene variants under strong selection, we identified regions of the genome with SNPs that were closely spaced enough that linkage could be established. Some regions with high densities of linked SNPs occurred in both the no-phage controls and post-phage populations (Fig. 6b) and thus were likely beneficial under all conditions (Fig. 6c). These included genes related to aerobic benzoyl-CoA catabolism, a type V secretion system, enoyl-CoA hydratase, and cell envelope structural proteins. However, there was a single region of the *Variovorax* genome that showed high SNP linkage density only in the post-phage populations. This locus encoded multiple conjugal transfer proteins (Fig. 6c). Most importantly, we identified two non-synonymous SNPs in the *trbE* conjugal transfer protein (homologous to *virB4*) within the P-loop domain which alters the charge and hydrophobicity of this region of the protein.

To identify SNPs that were not close enough to other mutations to provide linkage, we recovered all non-synonymous SNPs that increased in abundance by at least 30% in the post-phage populations relative to the pre-phage and no-phage control populations. We identified 275 gene variants found exclusively in post-phage populations (Supplementary data). Several were associated with known phage defence systems including the retron XIII, Lamassu family, and putative PDC-S02 defence systems. Most interesting were three SNPs detected in three or more of the four communities. This group of genes included a filamentous haemagglutinin family outer membrane protein, a redoxin/thioredoxin-like sensor protein associated with a fatty acid biosynthesis operon, and a protein with domain of unknown function (DUF932). This DUF932 gene occurs within an integrated plasmid that encodes *parA*, *parB*, an integrase, a plasmid transfer protein (*tral*), and a replication initiation protein (*repA*). Notably, multiple non-synonymous SNPs occurred in the DUF932 protein, and these variants were exceptionally abundant, exceeding 70% in the post-phage populations. In contrast, the *trbE* gene displayed non-synonymous SNPs in 21-33% of reads from post-phage populations. Interestingly, the start codon of this DUF932 gene was mutated to AAG (lysine) in 50-60% of each of the four populations. This likely impacts translation initiation and results in truncation of the protein as the next in-frame AUG codon occurs in position 66.

### Variovorax relies on its own thiamine biosynthesis in all enrichment cultures

To investigate genes important for *Variovorax* fitness in enrichment cultures (in the absence of phage), we introduced the *Variovorax* SCN45 RB-TnSeq mutant library into each of the four enrichment cultures (Fig. 7a). In all four of the communities, knockouts of multiple genes involved in the biosynthesis thiamine (*thiC*, *thiG*, *thiE, purE* and *purK)* had a strong negative fitness impact on *Variovorax* (See Fig. S6 for an overview of the pathway). These results indicate that in the four enrichments, *Variovorax* was reliant on its own biosynthesis for thiamine and little extracellular thiamine was likely available.

**Figure 7.**
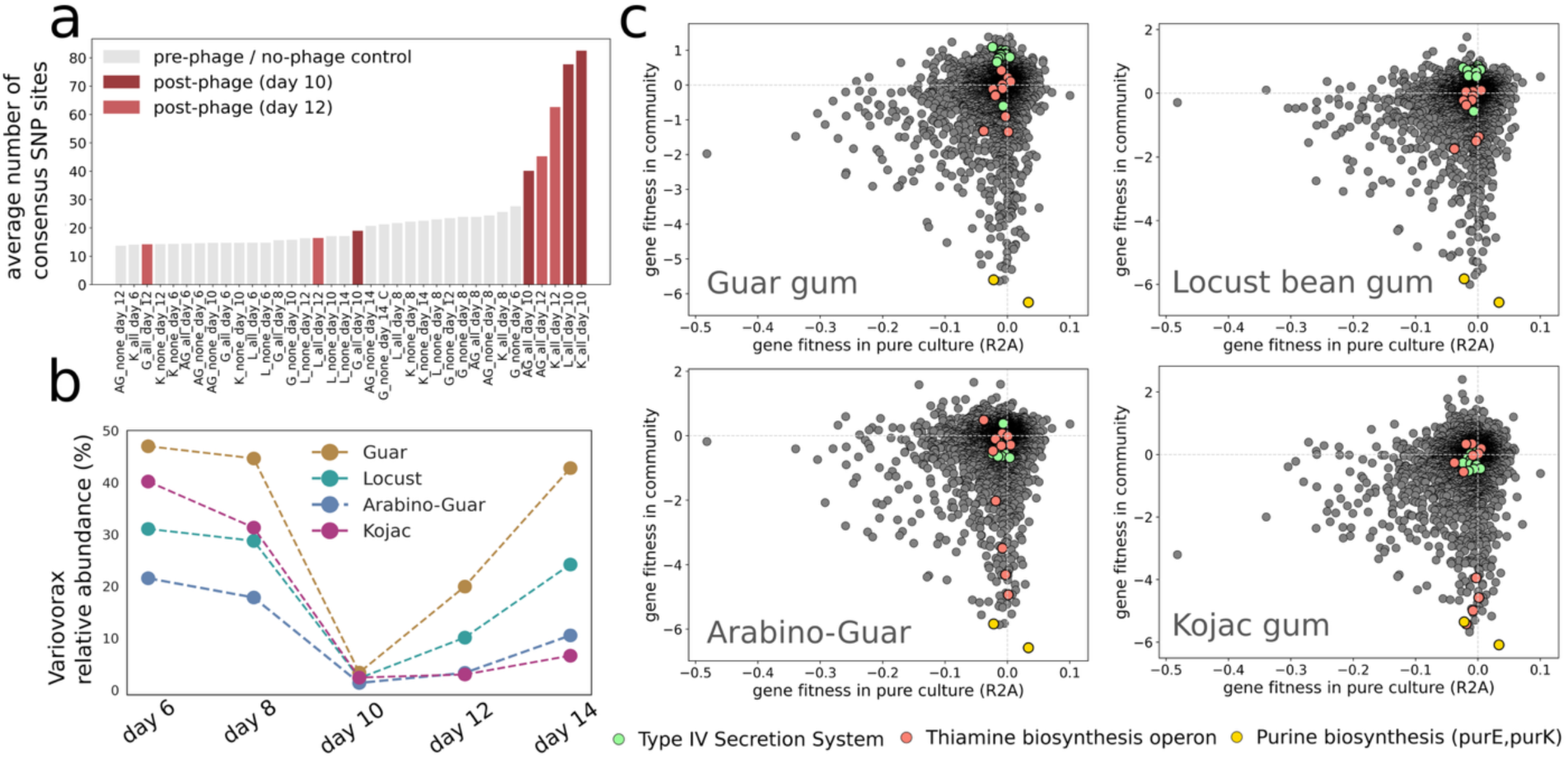
The rate recovery of *Variovorax* populations from phage infection corresponds to the fitness conferred by the type IV secretion system (T4SS). (a) Genome-wide fitness profiling (RB-TnSeq) for *V. SCN45* showing the fitness of gene-knockouts in each of the four communities (y-axis) versus in pure culture in R2A, rich media (x-axis), highlighting the type IV secretion system and thiamine biosynthesis genes. (b) Relative abundance of *Variovorax* is reduced to ∼3% in all post-phage communities, with a gradual recovery over the following passages, again fastest in the guar and locust bean gum communities. (c) Genome-wide SNP sites in *Variovorax* populations increase immediately following phage infection (day 10) but decline to pre-phage levels, with the guar and locust bean gum populations recovering the fastest. Read mapping was normalized to 20x coverage for all *Variovorax* populations.

### Rate of recovery of *Variovorax*populations correlates with reduced dependence on Type IV secretion system

Phage treatment reduced the abundance of *Variovorax* to approximately 3% in all day 10 post-phage communities, but *Variovorax* showed the fastest recovery in the guar and locust-bean gum communities over the following days (Fig. 7b). The *Variovorax* populations in these two communities (Fig. 7c, S5) had SNP concentrations close to pre-phage levels by day 14.

We amended enrichments with the RB-TnSeq mutant library to investigate the consequences of specific gene knockouts for *Variovorax* fitness in the four communities (in the absence of phages; Fig. 7a). In the guar and locust-bean gum communities the knockout of the T4SS was advantageous, based on selection for these genotypes. In contrast, these knockouts were mildly deleterious in the arabino-guar and kojac gum substrate-amended communities.

## Discussion

Previously, networks constructed using organism abundance data allowed us to identify thiamine auxotrophs that were strongly correlated with *Variovorax* (SCN45) in the bioreactor communities studied here (Hessler et al., 2023). Subsequently, we showed that *Variovorax* produces high concentrations of extracellular thiamine in pure culture, however we did not verify that this thiamine production was sustained within natural communities. To better explore possible interdependence related to vitamin provision required experiments conducted in communities from which *Variovorax* could be selectively removed, without directly impacting the rest of the community. We hypothesized that highly targeted community manipulation could be achieved via use of *Variovorax*-specific phages.

To demonstrate the feasibility of testing organism-organism interactions using phage-mediated *Variovorax* depletion, we leveraged the *Arabidopsis*-*Variovorax* CL14-*Arthrobacter* system first characterized by Finkel et al. (2020) in which *V. CL14* mitigates root stunting in *Arabidopsis* by degrading auxin produced by *Arthrobacter*. Following the addition of isolated phages V.cl_11 and V.cl_23 to this co-culture, we observed a pronounced root-stunting phenotype, indicating that the phages successfully depleted *Variovorax* CL14 (Fig. 2). These findings highlight the potential of extending phage-mediated testing of microbe-microbe and microorganism-plant interactions to rhizosphere microbiome manipulations for agricultural applications.

Next, we used three isolated *Variovorax* SCN45 phages to selectively deplete this *Variovorax* from four enrichment communities and monitored the effects on the remaining community (Fig. 4, 5). The subsequent addition of thiamine facilitated the recovery of only a *Leifsonia*, a Xanthomondales, and a *Microbacterium*, each lacking complete thiamine biosynthesis pathway. This provides clear evidence that *Variovorax*-produced thiamine was supporting the growth of these specific organisms in the communities and demonstrates the effectiveness of phage-mediated knock-down experiments for capturing these metabolic interactions. Related, but non-identical, strains of *Leifsonia*, *Microbacterium,* and the Xanthomondales were among a larger group of organisms associated with *Variovorax* in the correlation network from which the thiamine-exchange hypothesis was developed (Hessler et al., 2023).

Using a *Variovorax* SCN45 RB-TnSeq mutant library we found that disruptions in the *Variovorax* type IV secretion system genes conferred phage immunity, indicating that the three phages tested all infect via this complex. The T4SS is a common target for phage attachment and genome injection (Russell 1995; Berry et al., 2011). One potential concern is that depletion of *Variovorax* might occur following phage infection, yet decreased abundance of correlated organisms could be due to toxicity following uptake of viral DNA via T4SSs. Countering this, we find that encoding of a type IV secretion system was not a predictor of whether an organism decreased in abundance following phage addition. Thus, we concluded that these phages could be used to directly identify the organisms that are dependent on *Variovorax* SCN45 in the bioreactor consortia and the basis of the dependence.

*Variovorax* populations demonstrated partial to complete recovery from phage infection by the end of the experiment. We were unable to identify any gene deletions, frameshift mutations or non-synonymous SNPs in T4SS genes. However, by investigating genes with enriched non-synonymous SNPs, we identified gene variants of the *TrbE* conjugal transfer protein in *Variovorax* that arose following phage treatment. TrbE physically interacts with the T4SS and is a member of the *VirB4* family of proteins responsible for substrate recruitment and transfer to the secretion channel of T4SS (Li et al., 2018; Redzej et al., 2022). We observed two non-synonymous SNPs within the P-loop domain of this protein which is responsible for ATP binding and hydrolysis. We speculate that the *trbE* mutations interfere with the functioning of the T4SS required for phage infection.

In addition to *trbE* variants, we observed non-synonymous SNPs in a small-protein-coding (DUF932) gene within a putative integrated plasmid in all phage-treated *Variovorax* populations. Although the precise role of this gene is unclear, its consistent selection after phage application strongly suggests involvement in phage defence. The blocking of phage entry by resident mobile genetic elements, including phage, is well-documented (Kliem & Dreiseikelmann, 1989; Berry and Christie, 2011; Kamruzzaman et al., 2022; Johnson et al., 2022; Taylor et al., 2024; Rivard et al., 2024;). For example, the *pVII* minor capsid protein in *Pseudomonas* phage *Pf4*, interacts directly with the host *PilC* protein, responsible for T4SS assembly in *Pseudomonas*, rendering it inactive and preventing phage superinfection (Wang et al., 2022). We suspect that *Variovorax* developed phage resistance through both chromosomal and mobile element-mediated modulation of the T4SS. These mutations are likely to have existed within the *Variovorax* populations prior to phage application and highlight the role of population variation in surviving phage infection.

The addition of a single or multiple phages showed little difference in *Variovorax* depletion or its rate of recovery. This absence of additive effects has been observed in other studies (Pyenson et al. 2024), including efforts to remove *Ralstonia* from tomato plants (Franco Ortega et al., 2024). In our study, this may have occurred to all three *Variovorax*-targeting phages infecting via the same receptor. In future studies, the use of multiple phages targeting different receptors would likely be beneficial, as has been shown in phage-therapy research treating bacterial infections in humans (Yoo et al., 2024). However, complete bacterial removal is rare and, in clinical cases, has required the simultaneous use of antibiotics (Pirnay et al., 2024). Notably, in the *Ralstonia*-infected tomato plant studied by Wang et al. (2019), the addition of phages resulted in the enrichment of taxa that naturally suppressed the growth of *Ralstonia* through antibiotic production. These studies suggest that phage application alone is unlikely to remove target bacterium long-term, and that additional ecological or therapeutic pressures are required to sustain suppression of the target organism. However, as we show here, even partial and temporary depletion of populations is useful to test interaction hypotheses.

*Variovorax* recovered at variable rates following phage treatment, with communities in which T4SS knockouts conferred positive fitness exhibiting the fastest recovery. Prior work has shown that community composition can substantially impact phage-host dynamics (Blazanin and Turner, 2021; Alseth et al. 2019). Phage susceptibility can also be strongly influenced by the natural variation in physiology of genetically identical cells, such as cell size (Schenk & Sieber, 2019) and receptor protein copy numbers (Chapman-McQuiston & Wu, 2008; Igler et al., 2022). This leads us to hypothesize that community– and environmental selection against T4SS expression in *Variovorax* in the guar and locust bean gum communities could have led to reduced presentation of this complex by *Variovorax*, thereby limiting infection by our phages targeting this system. The observed variation in *Variovorax* recovery highlights how community context could impact phage infection, and may be an important area of research for the application of phage-based microbiome manipulations.

Here, we establish the basis of specific microbe-microbe interactions in natural enrichment-based communities by isolating and using phages to selectively deplete a member of interest. This approach addresses a critical challenge in microbial ecology by easily enabling the direct testing of interaction hypotheses within natural consortia, including interactions involving taxa that are otherwise difficult to culture or manipulate. This method is scalable, cost-effective and uses standard microbiology techniques without requiring specialized equipment. Although suitable phage isolation does require the isolation of the microorganism of interest, future integration of phage engineering with advances in protein design (Pacesa et al., 2025) may circumvent this current limitation. Overall, phage-based approaches represent valuable tools for understanding the structure of microbial communities and offer promising avenues for microbiome engineering.

## Methodology

### Bacterial inoculum preparation

*Variovorax* was grown overnight from a single colony in R2A media. 100 mLof planktonic phase cells were recovered from two bench-scale thiocyanate-degrading bioreactor systems, from which *Variovorax* was initially isolated a year prior. Flocculated particles were allowed to settle to the bottom of the container for 30 min and the top 50 mLwere carefully removed. 10 mLof bioreactor planktonic phase were added into 50 mLfalcon tubes. 5 mLof *Variovorax* overnight culture was added to this falcon tube. 15 mLof 50% glycerol was then added and mixed by swirling and inverting each tube. 200 uL of each culture were distributed into cryo tubes, frozen in liquid nitrogen, and stored at –80C. This approach was based on previous inoculum ratio testing experiments (Gao et al., 2021), which demonstrated that a higher abundance of a member in the inoculum improves its chances of establishing within the community.

### Community passaging

Inoculum glycerol stocks were thawed on ice and gently mixed by pipetting. 50 uL of glycerol stock was added to 10 mLof M9 minimal media in quadruplicate supplemented with 50 ug/L cycloheximide, 10 mg/L pantothenic acid, and a trace metal solution described by Watts et al. (2019). Culturing was performed using four independent technical replicates to provide robustness for statistical tests. The electron donors used were 0.2 g/L of kojac gum, locust bean gum, guar gum, and combined 0.1 g/L guar gum and 0.1 g/L arabinogalactan. Under each electron donor condition, following inoculation, cultures were incubated at 30C for 48 hours with shaking at 140 rpm. Cultures were passaged using 200 uL of spent medium into 10 mLof fresh media. This was repeated every 48 hours for a total of seven passages over 14 days. This culturing protocol was heavily informed by Goldford et al. (2018). The glycerol stocks were then used to inoculate fresh tubes followed by two passages as described above. At each passage step, 8 mLof culture were centrifuged at 10,000 X g for 10 minutes and cell pellets were frozen at –80C for subsequent DNA extraction. 200 uL of spent medium was used to perform OD measurements using a 96-well plate reader at 600 nm.

### Virome preparations

Three sources of viromes were collected: (i) soils, (ii) composts and (iii) and thiocyanate-degrading bioreactors. Briefly, approximately 10 grams of soil or compost were collected in falcon tubes and incubated with 25 mLof SM buffer with shaking for one hour. In the case of bioreactor viromes, biofilm material was removed from the bioreactors using serological pipettes and subjected to metal bead beating on a vortex at maximum speed for ten minutes to disrupt the biofilm. For all three types of viromes, the particulate matter was then allowed to settle for 30 minutes before the supernatant was removed and centrifuged at 10,000 X g for 10 minutes and the supernatant then passed through a 0.22 um filter.

### Phage liquid enrichment

Bacteria were grown overnight from single colonies in 10 mLof R2A media and incubated at 30C at 160 rpm. Cultures were back diluted into fresh R2A containing 2 mM MgCl2 and 2 mM CaCl2 to an OD600 of 0.03-0.05. Cultures were allowed to grow until reaching an OD600 of 0.15-0.3. Subsequently, 100 uL of viromes were then added to the tubes and the cultures were incubated for a further 24 hours. The cultures were then sampled and cells were removed by centrifugation at 10,000 X g for 10 minutes. This supernatant served as the virome for the next round of enrichment. This process was repeated for a total of five enrichments.

### Phage isolation: Plaque assays

Plaque assays were performed on the *Variovorax* strains *V.* SCN45, *V.* CL14, *V. boronicumulans* (DSM 21722), *V. ginsengisoli* DSM 25185 and *V. paradoxus* DSM 645. These strains were grown overnight from single colonies in 10 mLof R2A media and incubated at 30C at 160 rpm. Cultures were back diluted into fresh R2A to an OD600 of 0.03-0.05. Cultures were allowed to grow until reaching an OD600 of 0.15 – 0.3. Subsequently, 100uL of log-phase cells were added to 20 uL of 0.2 um filter-sterilized viromes and incubated at RT for 10 minutes. These were mixed with 50C R2A (2 mM MgCl2 and 2 mM CaCl2) 0.5% top agar and plated onto R2A 1.5% bottom agar plates. Plates were incubated at 30C for 48 hours and inspected for plaques. Plaques were picked using sterile 100 uL tips into 100 uL of SM buffer.

### Phage genome sequencing, assembly and annotation

DNA from phage suspensions were extracted using Zymo Quick-DNA viral kits using manufacturers instructions. Extracted DNA was then Illumina shotgun sequenced on a Novaseq using 150PE reads with a 300 bp insert size and yielding 1 Gbp per sample. Viral genomes were assembled using Metaviral-Spades (Antipov et al., 2020) and extended using COBRA (Chen et al., 2024). The genomes were then annotated using vogDB (Trgovec-Greif et al., 2024), Pfam (Bateman et al., 2004) and KEGG (Kanehisa & Goto, 2000) hidden Markov models using hmmscan. Protein sharing of the nine phage and Refseq and was performed using Vcontact2 (Bin-Jang et al., 2019). Phage genome ANI was determined using Vclust (Zielezinski et al., 2024; https://github.com/refresh-bio/vclust).

### Transmission electron microscopy (TEM)

Virus particles were visualized using a FEI Tecnai T12 Transmission Electron Microscope at an accelerating voltage of 80 kV. Carbon-coated copper grids were charged in a vacuum chamber followed by application of 5 uL of virus suspension and allowed to dry. The grids were washed with deionised water three times before negative staining in 1.0 % ammonium acetate for 60s. Grids were allowed to air dry before being visualized.

### *Arabidopsis* inoculation and co-cultures with *Variovorax* CL14, *Arthrobacter* CL28, and V.cl_11+V.cl_23 phage for root length phenotyping

*Arabidopsis thaliana* Col-0 seeds were surface-sterilized in 50% bleach for 20 min with 0.05% Tween 20 for 20 min, removing all bleach and rinsing five times in sterile water. Sterile seeds were stratified in the dark for 2 to 3 days at 4°C. A nylon mesh filter with 100 micron pore size was cut to fit on the 100 mm x 100 mm square petri dish as previously described (Cole et al. 2017). After stratification, ∼7 seeds were placed on the mesh filter on 1.0MS, 0.5% sucrose, 0.6% phytagel plates and grown under long day conditions for 7d (16-h light/8-h dark regime at 22-°C). Bacteria strains preparation: *Variovorax* CL14 and the root growth inhibiting strain *Arthrobacter* CL28 were streaked on agar plates from glycerol stocks, a single colony was transferred to 3 mL liquid R2A media and grown overnight. Cultures were gently pelleted then washed twice with 10 mM MgCl2, normalizing the volume of each culture to OD600 = 1, then adjust to OD600=0.04 before spreading 50 μl on each plate (Full MS, 0.6% phytagel without sucrose. Full MS was used instead of 1/2X as previously described to maintain the required Mg and Ca concentrations for optimal phage lysis (2mM)). To prepare the phage inoculation, 5 ul of a phage cocktail (Siphoviridae phage V.cl_11+V.cl_23; 1×10^4^ PFU/uL each) was diluted in 500 uL 20 mM MgCl2 and 100 ul of this solution was spread onto each plate. After the plants grew 7d, the mesh was transferred to each of the bacteria and phage inoculated plates and returned to the growth chamber vertically grown in short day conditions (16-h dark/8-h light regime at 22°C). Plates were imaged 10 days after transfer. Primary root length was measured using ImageJ software, first setting the scale to the grid on the plate (13 mm) before measuring primary root elongation for each plant. There were three replicate plates (containing n≥5 plants each) for each control and treatment: uninoculated, no bacteria + V11/V23 phage, *Variovorax* CL14, *Arthrobacter* CL28, *Variovorax* CL14 + *Arthrobacter* CL28, *Variovorax* CL14 + *Arthrobacter* CL28 + V11/V23 phage.

### Phage-perturbed community experiments

Stable communities were grown on selected carbon sources and exposed to *Variovorax*-specific phages (V45_v66, or a three-phage cocktail of V45_v66, V.45_iii, and V.45_bapic4) 8 hours after the passaging of the day 8 communities (ie. following passage 5), using 50 uL of 4 × 10⁴ PFU/uL each per 10 mLculture. Experiments were conducted in quadruplicate with a no-phage control included for comparison. The final varied condition involved the presence or absence of 10 mg/L thiamine, administered at the time of phage addition and maintained during subsequent passages.

### DNA extraction, amplicon and shotgun sequencing

Genomic DNA was extracted from cell pellets using the Qiagen DNeasy PowerSoil Pro Kit according to the manufacturer’s instructions. V4-V5 16S rRNA gene amplicons were generated by PCR using primers 515F (GTGNCAGCMGCCGCGGTAA) and 926R (CCGYCAATTYMTTTRAGTTT) and cycling conditions 95C 10 min, 72C 90s, 50C 60s, 72C 90s for 30 cycles and a final extension at 72C for 10 min. PCR products were cleaned using Ampure XP magnetic beads. PCR products were sequenced by Innovative Genomic Institute’s Next Generation Sequencing Core on an Illumina MiSeq instrument using a read length of 2 × 150 bp with approximately 50,000 reads per sample. Generated amplicon sequencing data was processed yielding ASVs as performed by Kust et al. (2023). Whole genome community library preparation and shotgun sequencing was performed by Novogene. Paired-end Illumina libraries were prepared from genomic DNA with an insert size of 350 bp and sequenced on an Illumina Novaseq producing approximately 6 Gbp of raw sequencing data per sample. The *Variovorax* isolate was subsequently sequenced using PacBio long-read sequencing at 60x coverage, to generate two circular chromosomes.

#### Metagenomic Read Processing, Assembly and binning

Illumina 150 PE reads from community shotgun sequencing were processed as documented in the ggKbase data preparation pipeline (https://ggkbase-help.berkeley.edu/overview/data-preparation-metagenome/). Reads were processed using BBtools (Bushnell 2020) and sickle (Joshi and Fass, 2011), and were then assembled using IDBA-UD (Peng et al., 2012). Open reading frames (ORFs) were predicted using Prodigal V2.6.3 (Hyatt et al., 2010). Contigs were binned using CONCOCT (Alneberg et al., 2013), Maxbin2 (Wu et al., 2016), MetaBat2 (Kang et al., 2019), and VAMB (Nissen et al., 2021). The best bins per sampled were selected using DAS tools (Sieber et al., 2018) and a final non-redundant genome set across all enrichment communities and the previously published thiocyanate bioreactor genome set found at https://ggkbase.berkeley.edu/SCN_92/organisms was selected using dRep (Olm et al., 2017) at 95% ANI.

### Microbial genome annotation, classification and strain analysis

Microbial genomes were then annotated using hmmscan against Pfam (Bateman et al., 2004) and KEGG (Kanehisa & Goto, 2000) databases and were taxonomically classified using GTDB-tk (Chaumeil et al., 2020). All read mappings were performed with bowtie2 (Langmead et al., 2012). Genome coverage and relative abundance across samples were determined using coverm (Aroney et al., 2025). Genome-genome correlation was determined using Fastspar (Watts et al., 2019) which implements the Sparcc algorithm (Friedman et al., 2012). *Variovorax* strain analysis was performed using inStrain (Olm et al., 2021) and the PacBio assembled *Variovorax* genome. Phage defence systems were predicted in the *Variovorax* genome using padlock (Payne et al., 2022, https://padloc.otago.ac.nz/padloc/). Protein structure predictions were performed using Alphafold3 (Abramson et al., 2024. https://alphafoldserver.com/)

### Statistical analyses

Distributions in organism abundances were compared by two-sample t-test assuming unequal variance was performed using the ttest_ind function of the scipy.stats Python package. The abundance of ASVs was calculated excluding the abundance of *Variovorax* contributing to 100% abundance. Relative abundance data were normalized by robust centered log-ratio (rclr) transformation using the decostand function from the vegan R package (Oksanen et al., 2001). Principal component analysis (PCA) was performed on the normalized abundance data using the prcomp function in R to visualize variation in community composition across samples.

### RB-TnSeq library construction

A DNA-barcoded *mariner* transposon mutant library of *Variovorax* SCN45 was generated as described previously (Wetmore et al., 2015; Thorgersen et al., 2021). Briefly, an *E. coli* strain (WM3064) carrying the transposon vector library pHLL250 was conjugated with wild-type *Variovorax* SCN45 at a 1:1 ratio on LB agar plates supplemented with diaminopimelic acid (300 uM) for 6 hours. Post conjugation, the transformants were plated onto R2A agar plates containing 50 μg/ml kanamycin. Several thousand *Variovorax* colonies were scraped and collected into a tube before being resuspended in 100 mL of R2A with Kanamycin and incubated at 30 C for 6 hours. Glycerol stocks (15%) were made and stored at –80C. Cells from the final mutant library were pelleted through centrifugation and their DNA extracted and sequenced by TnSeq-like Illumina sequencing protocol.

### RB-TnSeq library incubations

A 2 mLmutant library glycerol stock was thawed and inoculated into 25 mLof R2A with 50 ug/ml kanamycin and incubated at 30°C with shaking until cells entered mid log phage (approximately 8 hours). Four tubes of 2 mLof culture, termed time zero controls, were then recovered and cells pelleted by centrifugation and frozen at –80°C. The remaining cultured cells were then washed twice in relevant media (M9 media without carbon source or R2A) with spins at 3000g for 3 minutes. The cells were resuspended in media and inoculated into relevant or culture to a final OD600 of 0.02. The cultures were grown in 10ml glass tubes and incubated at 30 °C with shaking. (i) For phage-challenge experiments 10 uL of phage (4×10^4^ PFU/ul) were added to the libraries after 4 hours of subsequent incubation and allowed to incubate for a total of 24 h. (ii) For community incubations, cultures were passaged every 48 h for four passages. Cells were recovered from four replicate cultures by centrifugation and stored at –80 °C. Genomic DNA was then extracted from the stored cell pellets and PCR and sequencing by BarSeq performed as described previously (Wetmore et al., 2015).

### Data availability

The contigs representing the *Variovorax* genome assembled by PacBio can be found under bioproject PRJNA1212206 genome accessions CP179718-CP179720. The assembled genomes of the isolated phage have been deposited in GenBank under accession numbers PV558788 – PV558796. The reads associated with these genomes and assemblies can be found under accession numbers SRR33449703 – SRR33449695. The 16S rRNA amplicon sequencing data can be found under Bioproject PRJNA1254325, under accession numbers SRR33264975 – SRR33264785. Shotgun sequencing can be found under bioproject PRJNA1254232, including reads (SRR33436150-SRR33436076), assemblies (SAMN48138949 – SAMN48139025), and novel MAGs (SAMN48122594 – SAMN48122609). These MAGs can also be found on ggkbase https://ggkbase.berkeley.edu/Phage_treated_Variovorax_enrichments_dereplicated_MAGs/ and MAGs detected in this study that had been previously published can be found on ggkbase https://ggkbase.berkeley.edu/SCN_92/.

## Acknowledgments

This material by m-CAFEs Microbial Community Analysis and Functional Evaluation in Soils, a Science Focus Area led by Lawrence Berkeley National Laboratory is based upon work supported by the U.S. Department of Energy, Office of Science, Office of Biological and Environmental Research under contract number DE-AC02-05CH11231. Thank you to the staff at the UC Berkeley Electron Microscope Laboratory for advice and assistance in electron microscopy sample preparation and data collection. We thank Jennifer M. Doudna for institutional support and supervision. Thanks to Leylen Miloslavich, Marta Ortega and Kaitlin Creamer for assistance with lab operations, Henrik V. Scheller and Benjamin J Cole for *Arabidopsis* seeds, and to Jeffery L. Dangl for the *Variovorax CL14* strain and Rebecca S Bart for the *Arthrobacter CL28* isolates.

## Supplementary materials

**Figure S1.**
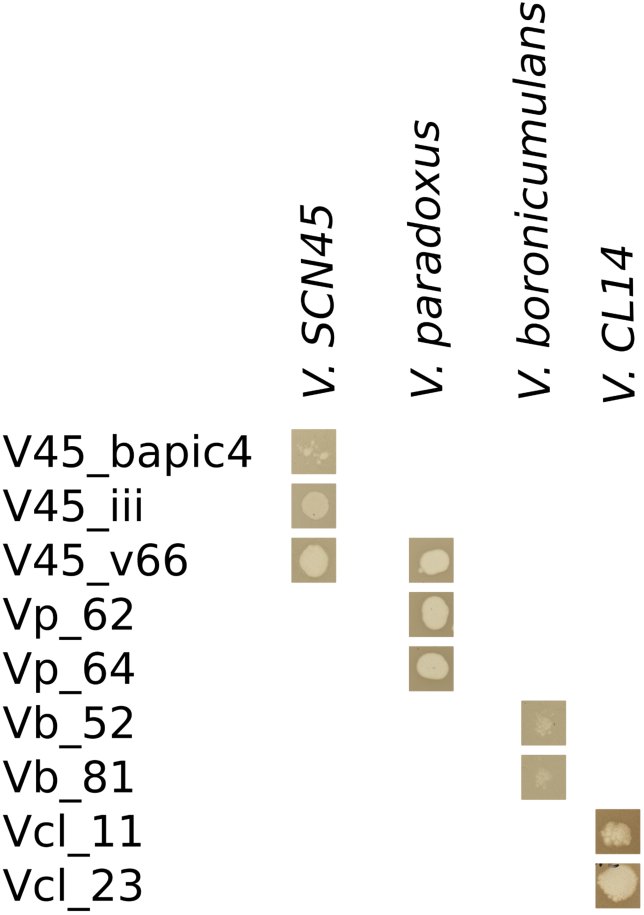
Plaque morphology from spot assays performed on *Variovorax* species with the nine isolated *Variovorax* phages. Only phage V45_66 was able to plaque on more than one host.

**Figure S2.**
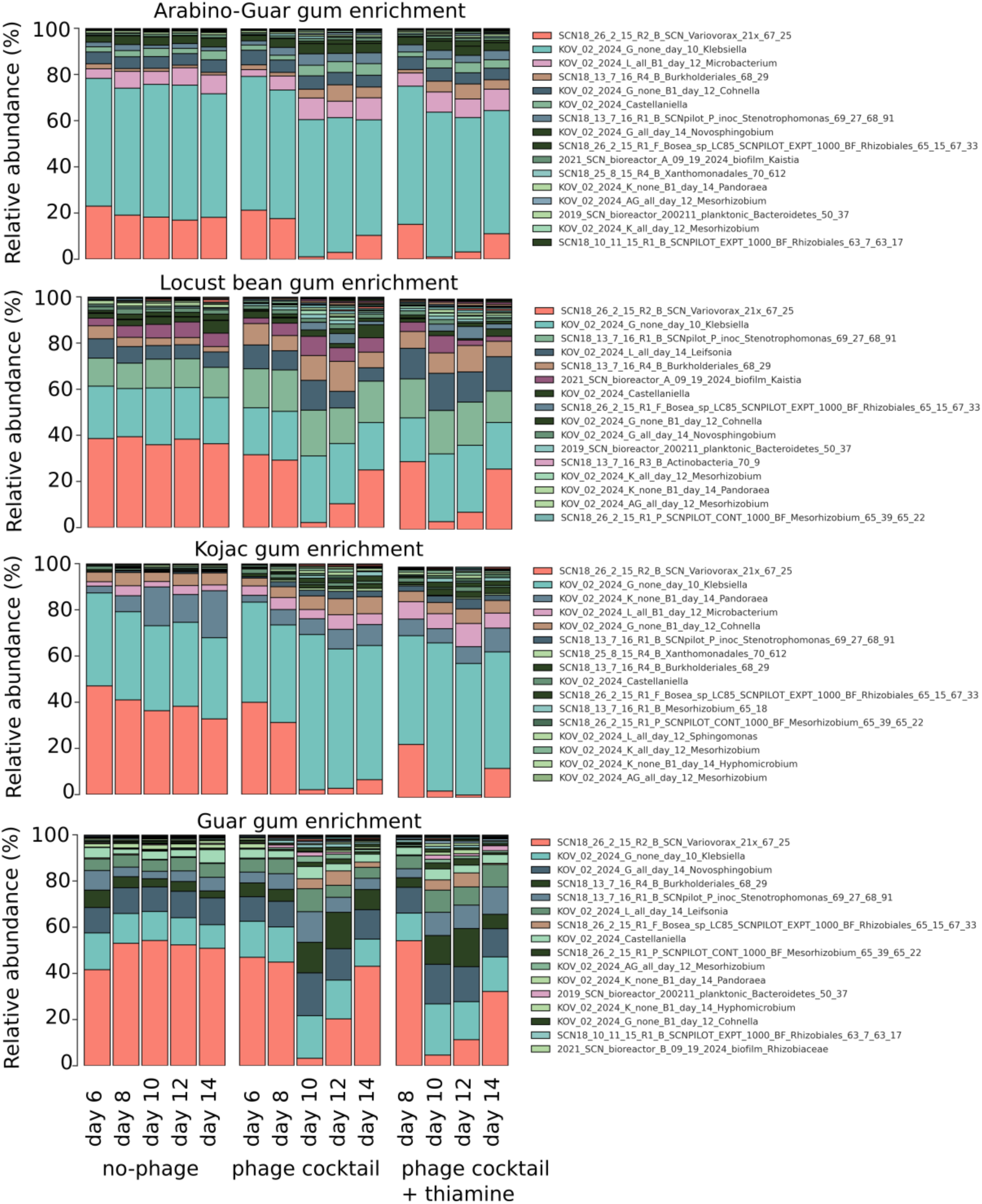
Genome-resolved metagenomics reveals consistent reduction in the relative abundance in Variovorax across all four enrichment cultures following phage application. The 16 most abundant organisms per enrichment culture recovered are shown.

**Figure S3.**
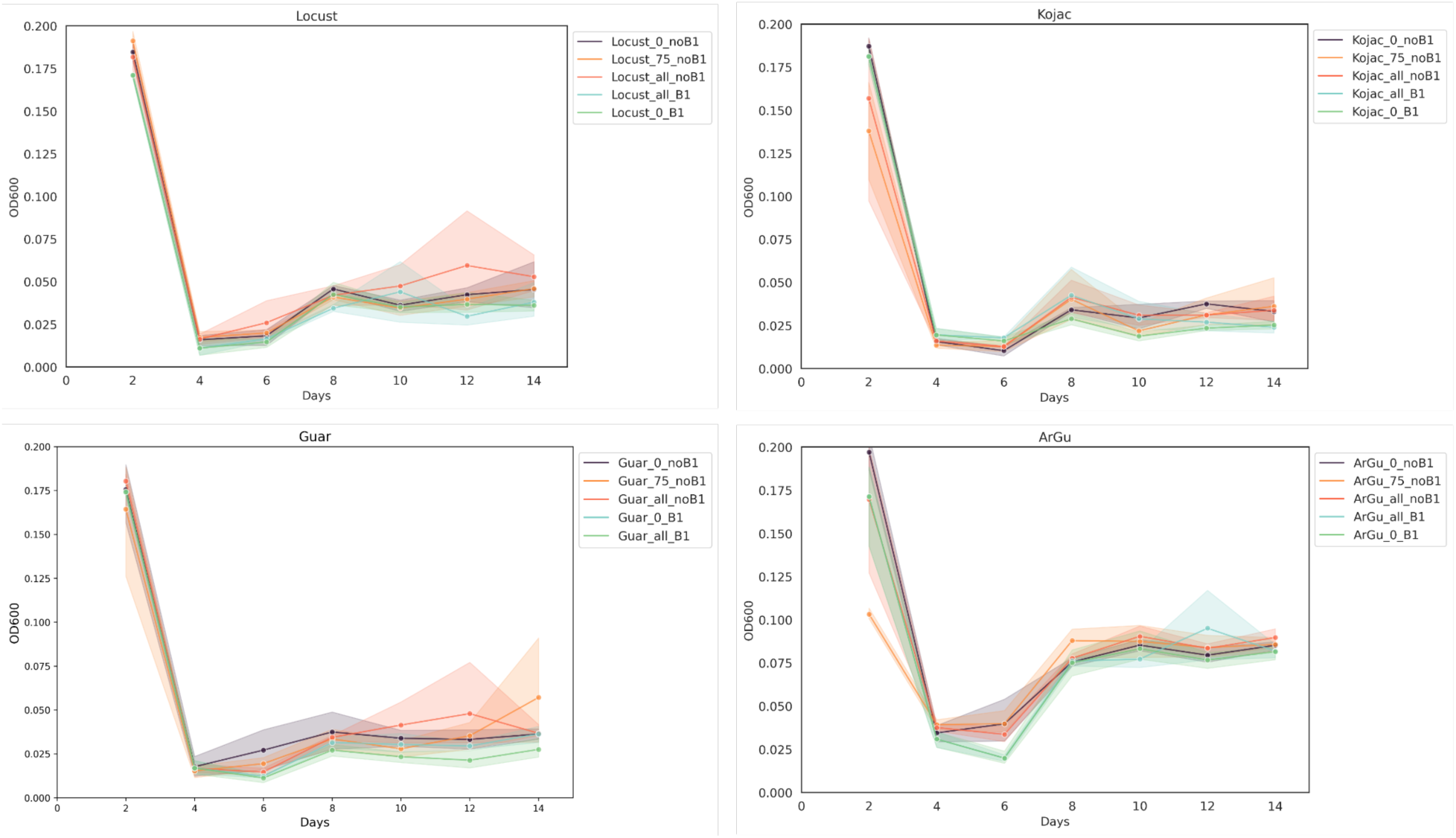
The optical density (600nm) of the 48 hour old communities following inoculation and each subsequent passage. All communities show a sharp reduction between day 2 and day 4 communities but then reach a stable state from day eight onwards.

**Figure S4.**
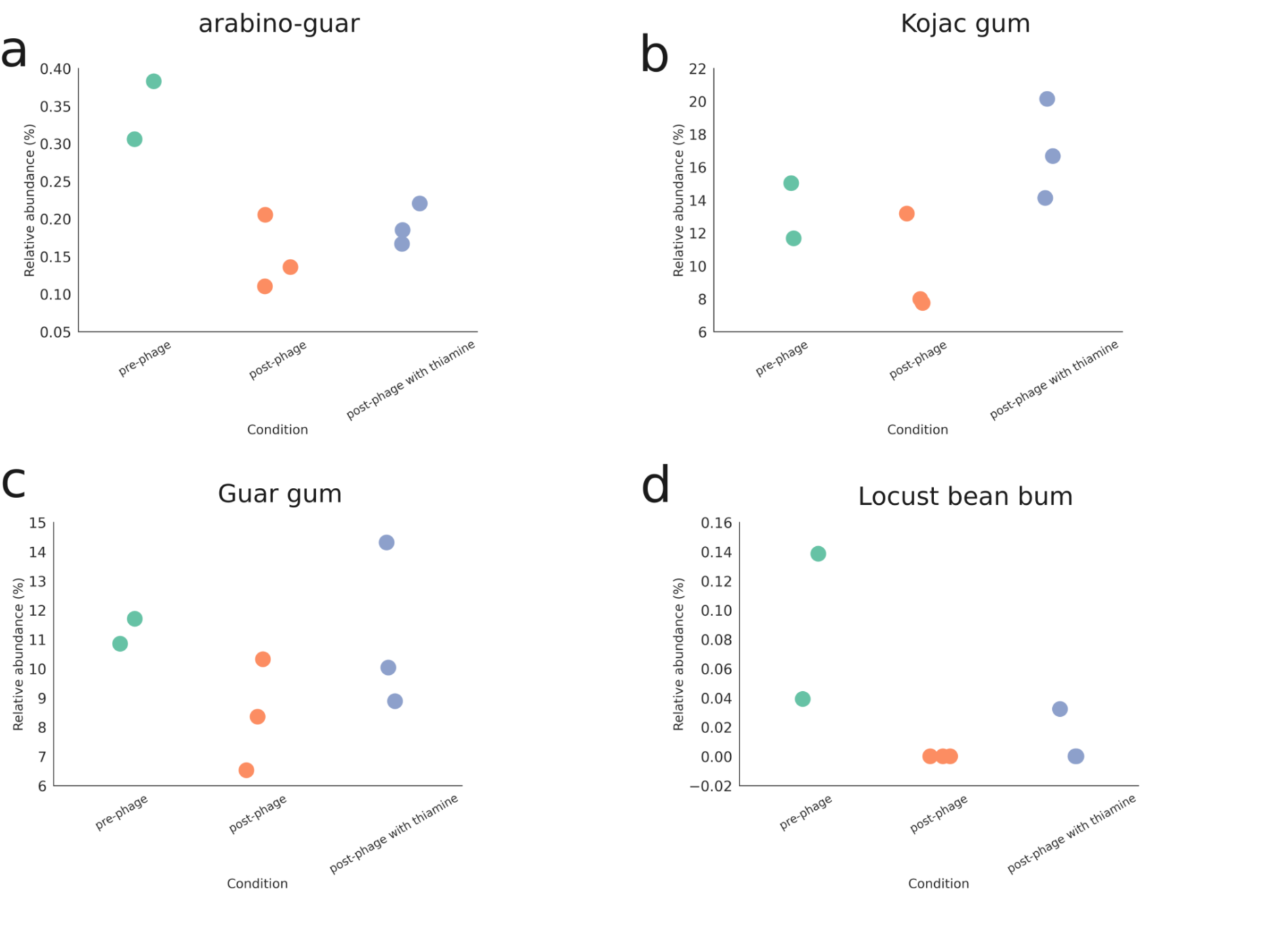
*Leifsonia* MAG relative abundance is consistent with the ASV data. The abundance of this MAG is shown across the four carbon source enrichment cultures before phage addition (day 6 and day 8 communities), following phage infection (day 10, day 12, day 14 communities), and following phage infection with supplementation of thiamine.

**Figure S5.**
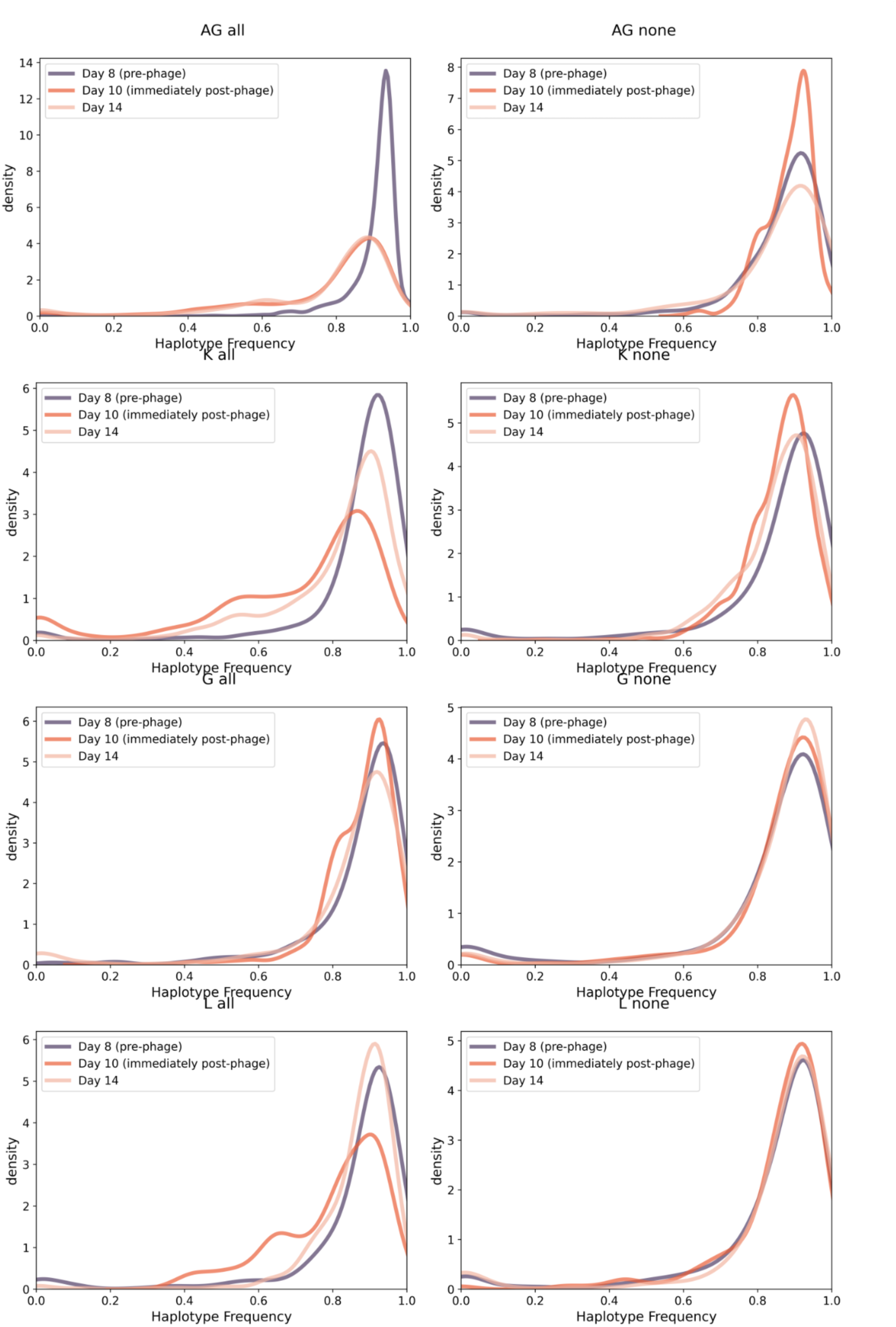
Haplotype frequencies associated with phage exposure to each of the four enrichment communities and their no-phage controls.

**Figure S6.**
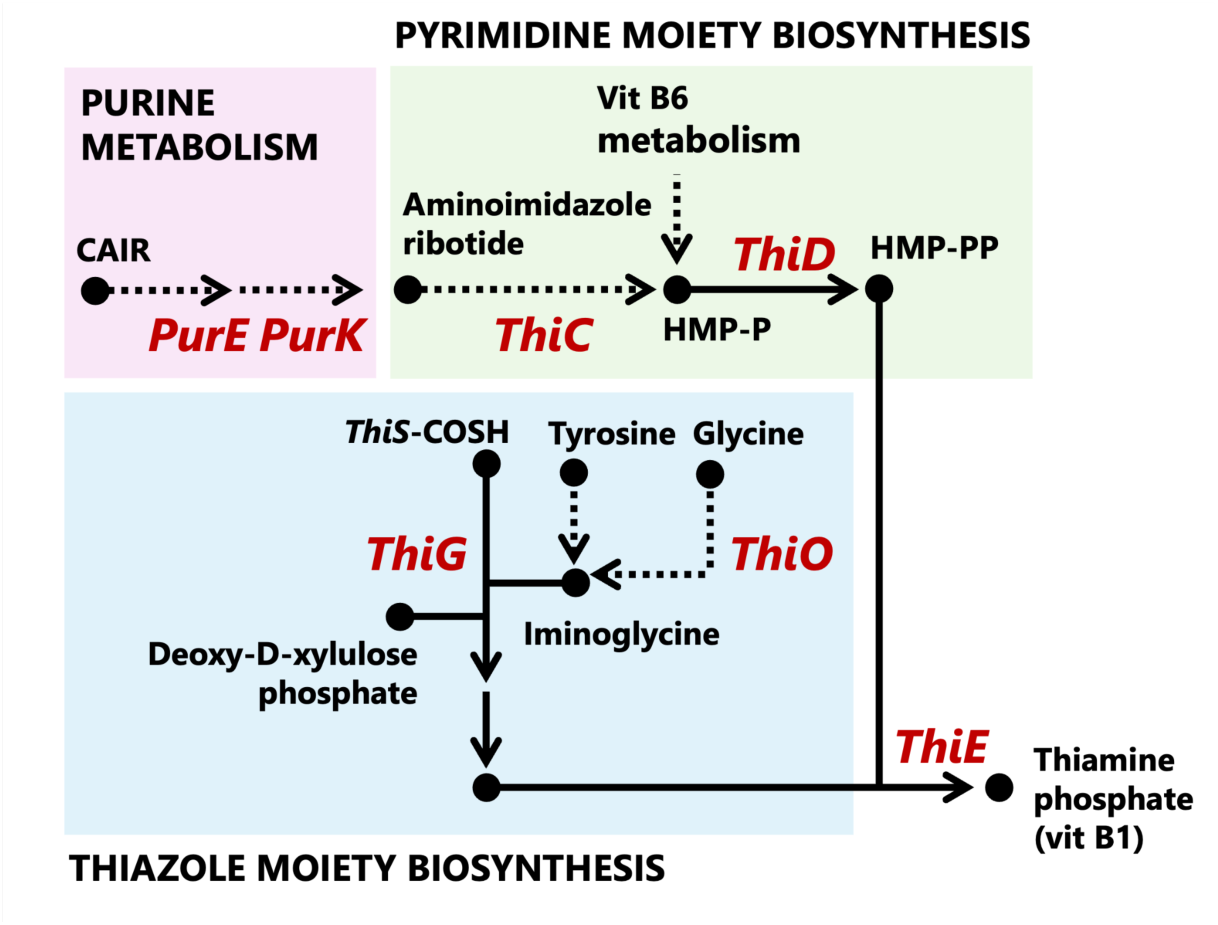
Thiamine biosynthesis pathway illustrating the multiple routes for the production of the thiazole and pyrimidine moieties of thiamine. Adapted from https://www.kegg.jp/ (Kanehisa & Goto, 2000).

